# New perspectives into the evolutionary pressures acting on the Human intrinsically disordered proteins

**DOI:** 10.1101/484659

**Authors:** Sergio Forcelloni, Andrea Giansanti

## Abstract

The codon usage bias is the well-known phenomenon of an unequal use of synonymous codons in coding DNA. These patterns reflect the action of weak selection working at the molecular level and allows to quantify the effects of natural selection, which tend to increase the fitness of the organisms. The prevailing hypothesis to explain the origin of codons usage bias is the *selection-mutation-drift theory*, according to which it results from a balance between the natural selection favoring optimal codons and combined action of random mutations and genetic drift which allow the persistence of nonoptimal codons. The main focus of this study is to quantify the extent of evolutionary pressures shaping the human genome. We found distinct patterns of mutational bias and natural selection in the human genes, depending on the structural properties of the encoded proteins (e.g. well-structured proteins, proteins with a long disordered segment embedded in a folded structure, or mostly unfolded proteins). Intrinsically disordered proteins are generally thought to evolve more rapidly, largely attributed to relaxed purifying selection due to the lack of structural constraint. Interestingly we observed that mostly unstructured proteins are not only affected by a basic mutational bias as the structured ones but are under a specific selective pressure underlining the important role of these proteins during evolution being freer to accept mutations, both neutral and selective. Our results provide new insight into understanding general laws and unknown aspects of protein evolution and they could be very useful in protein search and design.

## Introduction

Genetic information carried by mRNA and then translated in the corresponding protein is composed by triplets called codons. Four bases (A, C, G, U) compose the mRNA sequence so that there are 4^3^= 64 possible codons that, other than the three stop codons (UAG, UAA, and UGA), have to code for only 20 proteinogenic amino acids. Because more than one codon can encode for the same amino acid, the genetic code is degenerate or redundant. Indeed, with the exception of methionine and tryptophan, the remaining amino acids are encoded by more than one codon. For instance, lysine is encoded by two codons, isoleucine by three, alanine by four, and serine by six. Codons coding for the same amino acid are known as ‘synonymous codons’. Although these synonymous codons are indistinguishable in the primary structure of a protein, in a wide range of organisms, they are not used randomly, but with different frequencies which may vary across different species, in various part of the genome, and even within different regions of the same gene. This phenomenon is known as Codon Usage Bias (CUB) and it is well-established in the literature (Clarke 1969, Ikemura 1985, Plotkin and Kudla 2011, Shabalina 2013, Hanson and Coller 2017). Nevertheless, while remarkable observations have emerged, a clear interpretation of the role of the codon bias is still to be established because of its extremely complex phenomenology (Tuller 2014). The differences observed in the choice of codons between species can be traced to different evolutionary forces (Ikemura 1981). The most accepted theory to explain the origin of this phenomenon is the se*lection-mutation-drift theory* (Bulmer 1991). Based on this, natural selection favors codons that increase the efficiency and/or accuracy of the translational process acting on specific regions in the genome, which require a closer monitoring (Roth 2012); for example, selection on synonymous sites has been linked to transcription, splicing, DNA and mRNA secondary structure and stability, and protein folding (Zhou et al. 2009). On the other hand, the cooperation of random mutations and genetic drift allows the persistence of non-optimal codons and generally acts on the whole genome of an organism (Roth 2012). In particular, natural selection acting at the codon level for translational optimization may be of two types: positive or negative/purifying. The former favors mutations that increase the fitness of the organism; the latter tends to eliminate deleterious changes that could compromise the fitness of organisms slowing down the evolutionary process. That is clearly reflected in highly expressed genes, where a strong bias leads to use a narrow subset of codons, which are translated more efficiently and accurately (Shabalina 2013), and are optimized by co-adaptation to the tRNA pool in the cell (Ikemura 1985, dos Reis 2004). On the other hand, the persistence of non-optimal codons in specific regions along the mRNA sequence causes long breaks of the ribosome at the beginning of the ORF and during translation, which are crucial for the correct allocation of the ribosome on the translation start site (Zhou 2010) and for the folding process of the nascent polypeptide chain towards the native state (Spencer 2012). On the contrary of natural selection, mutational bias is almost neutral and does not substantially change the fitness of the organisms. Upstream are two main mechanisms by which neutral mutations can be generated. The first is related to the fact that the genetic code has degenerated. This is why many mutations can change a codon, but turn it into one that differently encodes the same amino acid; in this way a gene product will be exactly identical to the initial one. The second reason is that even when a mutation leads to the insertion of a different amino acid in the resulting protein chain, the functionality of the protein may not be affected; this, for example, if the substituted amino acid has the same chemical characteristics as the original one or if it is present in a site which is not essential for the purpose of the protein. These neutral mutations can be caused by different mechanisms that favor only certain types of mutations (e.g. non-uniform DNA repair, the chemical decay of nucleotide bases, and non-random error in the phase of replication).

The main difficulty in the analysis of codon usage bias in human is distinguishing the action of the natural selection from the neutral/mutational evolutionary processes. We remember that the mutational pressure is produced by distinct probability of different substitution types (Belalov 2013). Actually, a well-known feature of genomes, in general, is nucleotide or dinucleotide compositional bias. In the human genome a predominant factor driving the codon usage is believed to be GC content, i.e. the summed relative abundance of guanine (G) and cytosine (C) nucleotides; just think that the human genome was described as a mosaic of alternating low and high GC content ‘isochores’, large region of DNA (>> 200 kilobases) with a high degree uniformity in guanine and cytosine (Bernardi et al. 1985). The investigation reported in the article written by Bernardi et al. shows that the compositional compartmentalization of the genome (in heavy or light isochores) of warm-blooded vertebrates (like human) largely dictates the base composition of genes and their codon usage and this compositional constraint predominates over other constraints. Additional factors that might contribute to mutational pressure are methylation of CpG dinucleotide to form 5-methylcytosine and subsequent deamination that results in C-T substitution (Kaufmann and Paules 1996), chemical decay of nucleotide bases (Kaufmann and Paules 1996) and many mutations originated by methylation, by errors in replication, recombination, DNA repair, and by transcription-associated mutational biases (TAMB) (Comeron 2004). For example, it may be due to DNA polymerase which adds incorrect nucleotides; this can generate a transversion if there is an exchange of a purine with a pyrimidine or vice versa; a transition if there is an exchange of a purine with another purine or a pyrimidine with another pyrimidine. Moreover, as predicted by population genetics theory, the signature of translational selection in less conspicuous in humans than in species with much larger N_e,_ so that obtaining reliable estimates of selection on codon usage has proved complicated and may be detected only using a very large set of sequences. Therefore, in order to distinguish the extent of the pure selective pressure acting on the genome of a complex organism, we have to consider all this kind of compositional trends and the strong influence of background composition (isochores) in the coding sequence excluding them or minimizing their contributions in the calculation of the selective pressure. In this context, the phenomenon of CUB is very important because it provides examples of weak selection working at the molecular level and allows to quantify individual (site-specific) and combined (global) effects of the various evolutionary pressures acting on the genome of organisms. The focus of this work is on shedding some light about the different evolutionary pressures acting in the human genome. In particular, we would like to comprehend if the different form of protein disorder (e.g. well-structured proteins, proteins with a long disordered segment embedded in a folded structure, or mostly unfolded proteins) are under the control of a more or less pressing natural selection. To this aim, we consider a fine-tuned, sequence-only, broad classification of human proteins based on the presence of long unstructured segments and the percentage of intrinsic disorder. Two parameters control our distinction: dr, the percentage of disordered residues, and Ld, the length of the longest disordered segment in the sequence. We distinguish: i) *ordered proteins* (ORD), Ld < 30 and dr < 10%; ii) *not disordered proteins* (NDPs), Ld < 30 and 10% ≤ dr < 30%; iii) *proteins with intrinsically disordered regions* (PDRs), Ld ≥ 30 and dr < 30%; iv) *intrinsically disordered proteins* (IDPs), Ld ≥ 30 and dr ≥ 30%; v) *proteins with fragmented disorder* (FRAGs), with Ld < 30 and dr ≥ 30%. PDRs have been considered in the general category of intrinsically disordered proteins for a long time. We showed that PDRs are closer to globular, ordered proteins (ORDs and NDPs) than to disordered ones (IDPs), both in amino acid composition and functionally (Deiana A., Forcelloni S., Porrello A., Giansanti A., pers. comm. DOI: https://doi.org/10.1101/446351). In view of what I have just said, the study of CUB has allowed us to further distinguish our variants of disorder (ORDs, NDPs, PDRs, IDPs, and FRAGs) based on the magnitude of natural selection and mutational bias acting on them. IDPs are generally thought to evolve more rapidly, largely attributed to relaxed purifying selection due to the lack of structural constraint (Brown et al 2011). Coherently with Afanasyeva et al. (Afanasyeva2018), using different genetic tools (ENC-plot (Wright 1990), Neutrality plot (Sueoka 1988), PR2-plot (Sueoka 1995)), we observed that mostly unstructured proteins (IDPs) are not only affected by a basic mutational bias as the other variants but are under a specific selective pressure pointing to the idea that IDPs are important during the evolution of proteins being more freer to accept mutations, both neutral and selective. This analysis could be important to understand general laws and unknown aspects of protein evolution and very useful to predict which kind of proteins tend to vary faster in time exploring in a wider way the sequence space and thus giving rise to new functions in the cell. Moreover, it could also suggest new strategies for protein search and design.

## Methods

### Data sources

Human Coding DNA Sequences (CDSs) were retrieved by Ensembl Genome Browser 94 (https://www.ensembl.org/index.html) (Zerbino et al 2018). Only genes with UniProtKB/SwissProt ID have been included to make sure we only consider coding sequences for proteins. Human proteome was downloaded from the UniProtKB/SwissProt database (manually annotated and reviewed section) (UniProt Consortium, 2015). We consider only CDSs that start with the start codon (AUG), end with a stop codon (UAG, UAA, or UGA), and have a multiple length of three. Each CDS was translated in the corresponding amino acid sequence and then we filter all sequences that do not have a complete correspondence with a protein sequence in UniProtKB/SwissProt (https://www.uniprot.org/uniprot/?query=reviewed:yes). Incomplete and duplicated gene, sequences with internal gaps, unidentified nucleotides were removed from the analysis. A list of 18214 human CDSs was generated; the coverage of the human proteome in SwissProt (reviewed) is 90%.

### Disorder prediction

Each residue in the sequences of human proteins has been classified either as ordered or disordered using MobiDB3.0 (http://mobidb.bio.unipd.it) (Piovesan et. 2018). We have used MobiDB because it is a consensus database that combines experimental and manually curated data (especially from X-ray crystallography, NMR, and cryo-EM), indirect sources of information on the disordered state of residues and disorder predictions for all UniProt entries using these tools: IUPred-short, IUPred-long, GlobPlot, DisEMBL-465, DisEMBL-HL, Espritz-DisProt, Espritz-NMR & Espritz-X-ray.

### Classification of disordered proteins in the human proteome

To partition the human proteome into variants of disorder we followed Deiana et al. (Deiana A., Forcelloni S., Porrello A., Giansanti A., pers. comm. DOI: https://doi.org/10.1101/446351). Two parameters control our classification: dr, the percentage of disordered residues in the whole sequence, and Ld, the length of the longest disordered domain in the sequence. We distinguish five variants of protein sequences, namely:

i. *Ordered proteins* (ORDs), that do not have disordered segments longer than 30 residues (Ld < 30) nor more than 10% of disordered residues (dr < 10%);
ii. *not disordered proteins* (NDPs), that do not have disordered segments longer than 30 residues (Ld < 30), with more than 10% but less than 30% of disordered residues (10% ≤ dr < 30%);
iii. *proteins with intrinsically disordered regions* (PDRs), that have at least one disordered domain longer than 30 residues (Ld ≥ 30) and are disordered in less than 30% of their residues (dr < 30%);
iv. proteins that are *intrinsically disordered* (IDPs), that have at least one disordered segment longer than 30 residues (Ld ≥ 30) and that are disordered in more than 30% of their residues (dr ≥ 30%);
v. *proteins with a fragmented disorder* (FRAGs), that do not have a disordered fragment longer than 30 residues (Ld < 30) and that, nevertheless, have at least 30% of their residues predicted as disordered (dr ≥ 30%).

Following this distinction, ORDs and NDPs are, clearly, proteins with a limited number of disordered residues and absence of disordered domains (Ld < 30). We considered ORDs in order to represent a variant of completely ordered proteins with dr < 10%. PDRs, unlike NDPs, are proteins characterized by the presence of disordered domains (Ld ≥ 30) in structures that may well be globally folded. IDPs intended to be the true intrinsically disordered proteins, which not only embed disordered domains but are also disordered in a relevant percentage of their residues. FRAGs, are a small number of proteins characterized by highly distributed disordered residues along the entire sequence.

### Codon bias indices

Different measures have been introduced to quantify the degree of codon bias for a given gene. Here we use the improved implementation of Effective Number of Codon (ENC) as defined in (Sun et al. 2012) and GC content, calculated for each gene and in all three positions of triplets.

### ENC (Effective Number of Codons)

ENC was introduced by Frank Wright in 1990 (Wright 1990) and provides an estimation of the number of different codons used in a given gene; it is easily computed from codon usage data because it does not require a set of reference genes and quantifies the tendency of a gene to use a preference set of codons without making any assumptions regarding the identity of optimal codons. ENC is inversely proportional to the extent of nonuniform codon usage and ranges from 20 (when just one codon is used for each amino acid) to 61 (when all codon are used for each amino acid along the sequence). Coherently with what stated by Wright and in order to avoid stochastic sampling effects, we decided to consider genes longer than 100 codons (Wright 1990). To calculate the ENC we followed the improved implementation by Sun et al. (Sun et al. 2012), who separate the codon table in families according to the number of codons that encode for the same amino acid. In this way, there are 2 amino acid codified by one codon, 9 by two, 1 by three, 5 by four and 3 by six codons. Then, given a sequence of interest, the calculation of ENC starts by quantifying Fα, defined for each family α of synonymous codons:

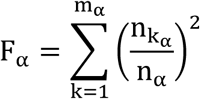

where m_α_ is the number of codons in the codon family α, n_iα_ with *i* = 1, 2, …, *m*_*α*_ is the number of occurrences of codon *i* of the codon family α in the coding sequence, and 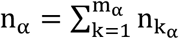. Finally, ENC weights these quantities on the sequence in the following way:

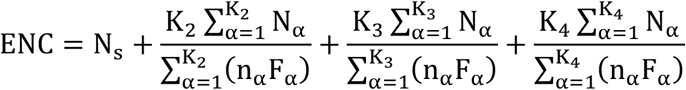

where N_s_ is the number of families with only one codon and K_m_ is the number of families with degeneracy m. In particular, given the structure of the genetic code, the families coded by 6 synonymous codons (Leu, Ser and Arg) can be split in two families, one with degeneracy 2 and another with degeneracy 4.

### GC-content

The GC-content of a sequence is the percentage of guanine or cytosine base respect to the total length of the gene. In the same way, it is possible to define the GC-content in the first, second and third position of codons that composed the sequence of a given gene. This measure is very interesting because it reflects the strength of the mutational bias on the genes. In the present analysis, the three stop codons (UAA, UAG, and UGA) and the codons that encode for the isoleucine were excluded from the calculation of GC3, and the two codons that code for Met (AUG) and Trp (UGG) were excluded from GC1, GC2, and GC3. Formally, the GC-content of a gene in a given codon position is defined by the following formula:

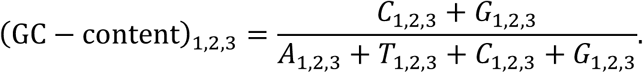

### ENC-plot

In genomic, the ENC-plot (Wright 1990) represents one of the tools used to study the equilibrium between the most important evolutionary pressures that determine the frequencies of the different codons: mutational bias and selective pressure. This graph is drawn using the ENC values as the ordinate and the GC3 values as the abscissa. As shown by Wright, a clear relationship between ENC and GC3 exists under the hypothesis of no selection, approximately expressed by the following formula:

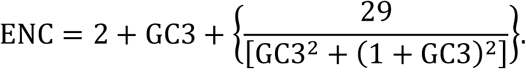

The meaning of the previous equation is clear. As we have already explained above, ENC quantifies CUB in a range from extreme bias (one codon used for each amino acid) to no bias (all codons are equally used for each amino acid along the sequence). Variations in GC3 make the estimation given by ENC more difficult to interpret. However, when there is no selective pressure that acts on the sequence the ENC value depends only on the effect of mutational bias quantifiable from the GC-content in the third codon position, which is the tendency of a sequence to put cytosine and guanine in the third position of each codon. In the extreme case, when no mutational bias (GC3=0.5) and no selection acts on the sequence, all 61 codons can be used and we obtain the explanation of the maximum value that we can observe in the curve defined by (5). Remaining in the situation of no selection, as soon as the mutational bias acts on the sequence causing some preference for codons that end with a certain type of bases (cytosine and guanine or timine and adenine), the codons are used unequally and some codon cannot appear in the sequence. In other words, stronger the mutational bias, lesser the used codons; it explains the tails and the decreasing trend of the curve for GC3<0.5 and GC3>0.5. As soon as the selective pressure acts on the sequence by influencing the use of codons, its corresponding point on the ENC-plot falls below the theoretical curve (5) and the distance between them provides us an estimation of the relative weight of the natural selection compared to mutational bias.

Before concluding, it is important to specify that we intentionally use the definition of ENC as defined by Wright (Wright 1990) and then revised by Sun et al. (Sun et al. 2012), not considering any correction for background nucleotide compositions (see for example Novembre 2002), as the aim of the first part is not to quantify the net extent of the codon usage bias but the balance between mutational bias and selective pressure that shapes the gene sequences leading to less uniform use of synonymous codons. Conversely, taking composition into account is particularly important for phylogenetic studies of codon usage bias where comparisons are made among species with differing background nucleotide compositions or for studies of codon usage bias in genomic regions that may differ in background nucleotide composition; however, this is not our case.

### Neutrality plot

The neutrality plot (Sueoka 1988) is another way to study the equilibrium of mutational bias and selective pressure. In synonymous codon usage, the base composition of the third position is the most vulnerable to mutational bias because the variations in that position in most cases do not change the corresponding amino acid. Because of the partially silent nature of the third position, the nucleotide in this position represents one of the most neutral within the genomes. Nevertheless, the neutrality of the third position does not mean that any change occurs in the third positions is neutral (Shabalina 2013). Conversely, change in first and in the second position are subjected to functional constraints because they may change the amino acid, except for some codons of Arg, Leu, and Ser. Following this reasoning, Sueoka considered the variation on the third position as an estimation of the extent of the neutral selection. So, studying the variation of GC1 and GC2 as a function of GC3, we can give an estimation of the entity of the equilibrium between mutational bias and selective pressure that constrains the first and second position of codons. Precisely, the neutrality plot is based on this idea and it is drawn using GC1 or GC2 on the ordinate and GC3 on the abscissa. The slope of the linear regression of GC1 and GC2 with GC3 expresses the equilibrium between neutrality and selective constraints. In particular, the slope is 0 for no effect of directional mutation pressure and 1 in the case of complete neutrality; it is worth to point out that stronger the mutational bias, more pronounced the slope of the linear regression coefficient. To be clearer, when the correlation between GC1 or GC2 and GC3 is statistically significant and the correlation coefficient is close to 1, the effect of the mutational neutral bias is dominant; conversely, when the correlation coefficient is near to 0, the effect of the selection (no-neutral) is dominant. The correlation analysis was carried out using the Spearman's rank correlation analysis method.

### PR2-plot

As already pointed out by Wright in his article, there are many cases where the codon usage data alone are insufficient to distinguish between mutational bias and selective pressure. If we observe the ENC-plot obtained for Homo Sapiens and D. melanogaster in (Wright 1990), in both cases the ENC value is highly correlated with GC3. However, in the first case this trend is due to variation in GC content of the mutational pressure acting in a different way in different regions of the genomes (due to the presence of large DNA segments that are characterized by an elevated GC-content, known as isochores), whereas in D. melanogaster, a similar trend is thought to be due to the effect of selection on the highly expressed genes which tend to have guanine or cytosine in the third position. The critical issue, therefore, is the relative roles of directional mutation pressure and selective codon usage bias on the variability of the DNA GC-content. To solve this ambiguity, we consider PR2-plot (Sueoka 1995). Soeuka, in his article, defined two types of intra-strand parity rules; the type 1 parity rule (PR1) is concerned with base substitution rates within one strand of DNA, and the type 2 parity rule (PR2) is concerned with the base composition at equilibrium within one strand of DNA. The principle PR2 and its violation in actual data provide a method to assess the relative contributions of directional mutation pressure, strand biases of mutation and selection to the base composition in terms of GC-content. To analyze and compare the PR2 violation, deviation from PR2 in genes of various organisms were examined for the third codon letter of four-degenerate synonymous codons (alanine, arginine4, glycine, leucine4, proline, serine4, threonine, and valine). If only neutral selection acts on the codon usage bias in a gene, that is in the absence of selection, it is expected that the base composition of a DNA strand will tend to equal frequencies of complementary bases among the 4-fold degenerate codon families (i.e. A=T and G=C in each strand), because changes in the third codon position of synonymous codons by substitution should not change the protein sequence. As soon as the selective pressure works on the sequence modeling its codon usage, its effect would lead to a violation of this proportion. Following this reasoning, the PR2 plot is a simple method to investigate the balance between the two types of pressures that influence the codon usage bias: neutral and natural selection. In particular, the PR2-plot is defined drawing the G3/(C3+G3)_|4-fold_ on the x-axis and A3/(A3+T3)_|4-fold_ on the y-axis, both of them measured on the 4-fold degenerate codon families. Because we are interested in analyzing the whole human proteome, we obtain wide spots of points (one point for each gene); if the neutral selection is the only one acting on the gene, the centroid of the distribution should be around the center where both coordinate values are 0.5; on the contrary, if the selective pressure influences the codon usage of the genes, the centroid of distribution moves away from the point (0.5,0.5). The distance of the centroid from the origin represents a measure of the strength of the selective pressure compared to the extent of mutational bias. In this way, PR2 rule and its violation provide a tool to complete the information obtained with ENC-plot.

### Excluding compositional bias from the calculation of the selective pressure

Unfortunately, the previous analysis based on ENC-, Neutrality-, and PR2-plot only allows us to understand if the various variants of disorder are subject to a different equilibrium between natural selection and mutational/compositional bias. However, they do not permit to specify the net extent of natural selection. In order to understand this aspect, we can make an analogy with a weighing scale with arms of equal length. This device does not allow to estimate the net weight of an object if the weight of the other object on the other arm is not known. In the same way, ENC-, Neutrality-, and PR2-plot represent different instruments (weighing scale) able to estimate the extent of one of the two selections (neutral or natural/Darwinian) in relation to the other. Moreover, ENC- and PR2-plot quantify the balance between natural selection and mutational bias measuring the departures from uniform distribution of synonymous codons; for our purpose, this is not a problem because the present work is a specialized (intra-species) analysis of the evolutionary pressures acting on the human genome and the aim of the first part is to investigate in which variant of disorder the effect of the mutational bias (including the bias in G+C content) is more pronounced. Having said that, the next step consists in characterizing the net extent of natural pressure acting on each variant of disorder. To do this, we have to take into account for the background nucleotide composition and filter all effects that can be derived from it. Novembre (Novembre 2002) developed a statistical correction (ENC’) that takes into account for background nucleotide composition. Although this is an improvement of the original method, ENC’ has its limitations: it is dependent from the length of the coding sequences, provides inaccurate estimates for short sequences (<600 bp) (Novembre 2002), and does not explicitly account for mononucleotide and dinucleotide composition bias that may be high in the human genomes (Belalov 2013). Moreover, it is possible that by using this method we eliminate some of the signal we’d expect to find. For these reasons in particular, we followed the procedure by Belalov et al. (Belalov 2013), who used different shuffling techniques of the coding sequences to quantify the net weight of selective pressure. All these techniques were designed to preserve the positional nucleotide and dinucleotide in the gene and the corresponding amino acid sequence, which is identical before and after shuffling. Python scripts were developed for dinucleotide content analysis and sequence shuffling (available on request). The algorithms of these scripts are reported in detail in (Belalov 2013) and are summarized below: i) *N*_3_ *correction* shuffled third positions of codons, thus precisely preserving genomic mononucleotide frequencies at the third codon position and the amino acid sequence; ii) *dN*_23_ and *dN*_31_ *corrections* shuffled dinucleotides between codon positions 2-3 (or 3-1, respectively), thus preserving genomic dinucleotide frequencies at the codon position 2-3 (or 3-1, respectively) and the amino acid sequence; iii) *dN*_231_ *correction* shuffled trinucleotides between codon positions 2-3-1 whilst preserving amino acid sequence.

## Results

### ENC-plot on the human proteome

To clarify the roles of the mutational bias and the selective pressure in determining codon usage bias within the different variants (ORDs, NDPs, PDRs, and IDPs) of proteins in which we have partitioned the human proteome and then, to estimate the magnitude of the codon usage bias within the human coding sequences, the ENC values were plotted against the GC-content at the 3rd codon position (GC3) for each gene in each variant of proteins. The plot (ENC vs. GC3s) was used to measure the codon usage of a gene that deviates from the equal usage of synonymous codons (Wright 1990). In Fig. 1 we show the ENC-plot obtained for the most representative variants of proteins: ORDs, NDPs, PDRs, and IDPs. From the curves fit on the bottom we can see that each variant has an ENC value below 61, which means that each variant uses a repertoire of codons different from the uniform distribution. All distributions are statistically different (p-value < 0.0001); on average, curves fit of PDRs and IDPs distributions are closer to the theoretical curve corresponding to the case when selection does not act on the coding sequences, meaning that neutral mutational bias affects more the codon usage bias of the variants containing at least one disordered segment longer than 30 residues than well-structured proteins, which are characterized by a low percentage of disordered residues in the sequence (ORDs and NDPs).

**Figure 1.**
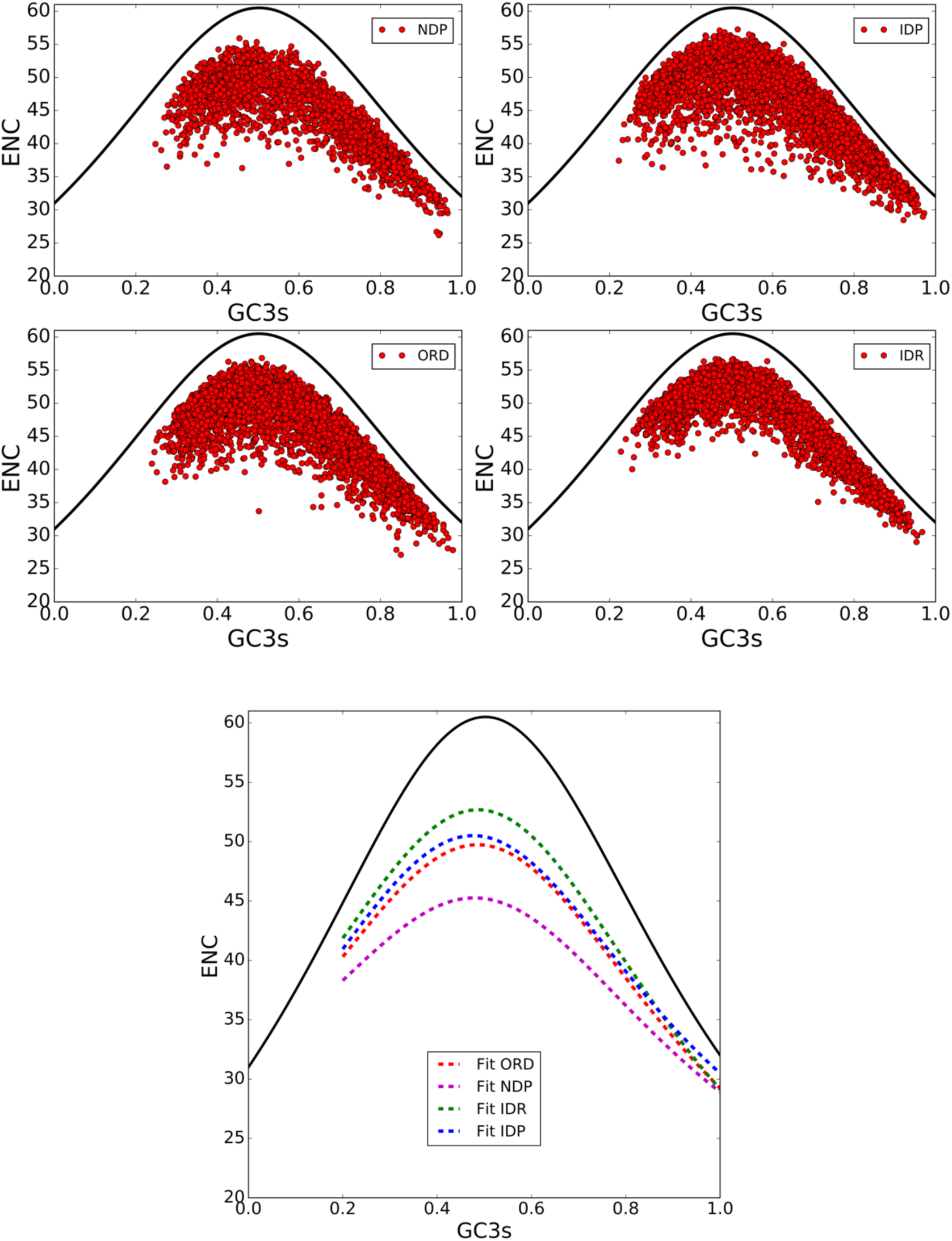
On the top, ENC-plot for each variant of proteins ORDs, NDPs, PDRs, and IDPs; on the bottom, we show the fit obtained parameterizing the theoretical curve suggested by Wright and defined in Methods.

### PR2 plot on the human proteome

As we have already specified in Methods, the ENC-plot analysis cannot completely distinguish between mutational bias and selection. To solve this ambiguity, we consider the PR2-plot. In Fig. 2 we report the PR2-plot on which we represent the various classes. Deviations from PR2 in A-T bases of the third position are shown as A3/(A3+T3)_|4-fold_ and those in C-G bases are shown as G3/(C3+G3)_|4-fold_ for the eight four-codon family amino acids. It is clear from Fig. 2 that the violation of PR2 is the rule rather than the exception, and the violation pattern is different for each class of proteins (ORDs, NDPs, IDRs, IDPs).

**Figure 2.**
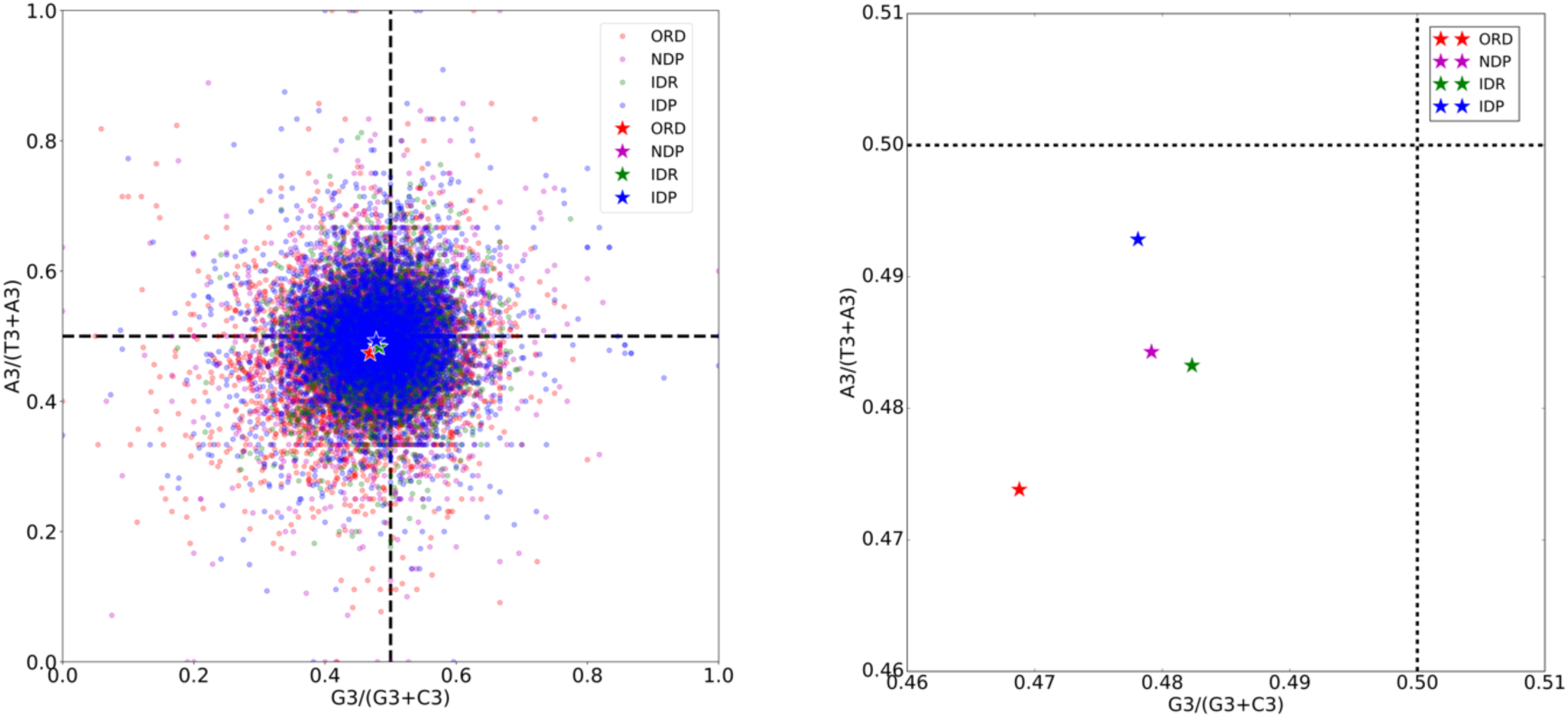
Representation of the classes ORDs, NDPs, PDRs, and IDPs on PR2-plot. The centroids of distributions are represented by stars with the same color as the points of the reference distribution.

To make this analysis clearer, for each point we calculate the distance 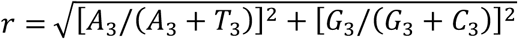 from the point of the coordinate (0.5,0.5) and, then, we construct the distribution of this variable for each variant. In Fig. 3 we show the distributions thus obtained; in addition, we represent the average values of the distributions with vertical colored lines with the same color as the points of the corresponding class in Fig. 2. The differences among the average values of the distribution observed in Fig. 3 are significant (Mann-Whitney test and p-value<0.001).

**Figure 3.**
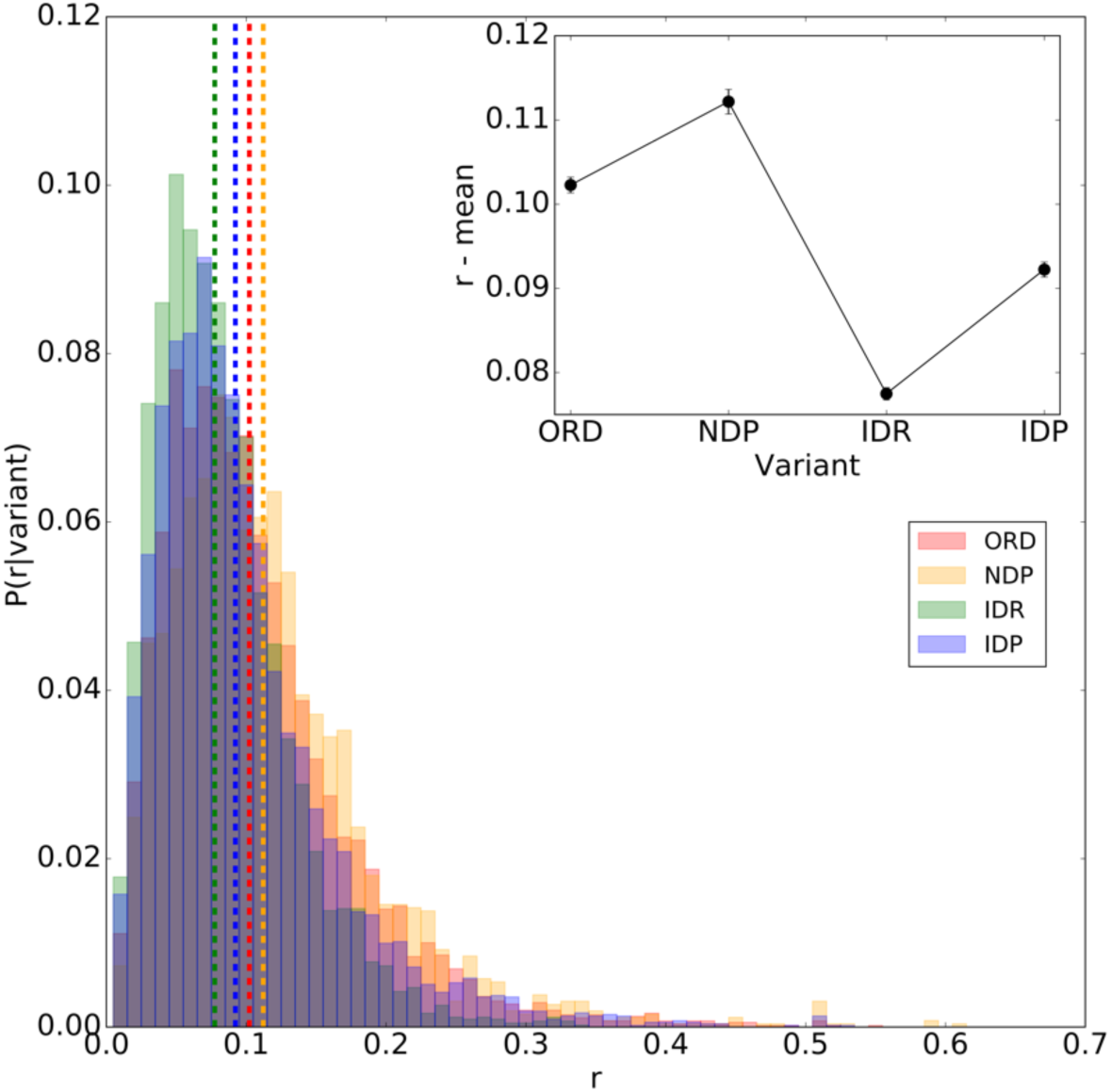
On the left, distribution of the distances 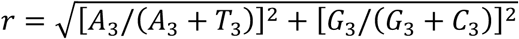 from origin in the PR2-plot. The vertical colored lines in the figure represent the average of the distributions. Starting from the origin (*r* = 0), the order of the variants is: IDRs, IDPs, ORDs, and NDPs. In the insert, we show the mean values of the distributions for each class of proteins.

Consistently with the ENC-plot analysis, centroids of PDRs and IDPs distributions are closer to the central point of the coordinate system (Fig. 3 and insert), meaning that neutral mutational bias affects more the codon usage bias of these disordered variants of proteins than ordered ones (ORDs and NDPs). In line with the ENC-plot analysis, completely ordered proteins (ORDs) and moderately disordered proteins (NDPs) move away from the central point. This observation is really interesting because it means that the net weight of natural selection compared to that of mutational bias in the ORDs and NDPs actually is lower than that of proteins containing at least one long disordered segment (PDRs) or a high percentage of disordered residues (IDPs). Consistently, the distances of centroids from the point (0.5,0.5), are perfectly coherent with the distances observed in ENC-plot between the curve fit of the variants of disorder and the theoretical curve corresponding to the case if no selection acts on the sequences.

### Neutrality plot and human proteome

It is well known that variations occurring in the third coding position are synonymous and almost weaker or neutrally selected; conversely, variations in the first and, especially, second codon positions are under the influence of a more marker pressure because they change the amino acid altering the phenotype. Having said that, the extent of natural selection could be estimated by the GC-content in the first and second positions, while mutational pressure could be mostly determined by the GC-content in the third codon position. In order to understand how the mutational neutral bias acts on the different positions of the codons, we used the neutrality plot (see Methods) which gives an idea of how the neutral bias acts on the first and second positions (in general no silent sites) compared to the third position (in general silent site). So, we performed separately a correlation analysis between GC-content in the first and second positions with the GC-content in the third position. Doing this, we construct the neutrality plot for each variant of proteins (ORDs, NDPs, PDRs, and IDPs); the slope of each linear regression gives an idea of the strength of natural selection compared to neutral selection. As we can see from the slopes of the linear regressions in Fig. 4, we found a significant but weak correlation which supported the influence of selective pressure on the codon usage bias of all variants of disorder. In particular, the ordered classes (ORDs and NDPs) seems to be more affected by natural selection with respect to the disordered ones (PDRs and IDPs), especially if we observe the large splitting among the slope in the GC2 vs GC3. It is worth noting that the slope in GC1 vs GC3 plot is greater than those obtained for GC2 vs GC3 plot. This observation is entirely reasonable because the second position is more under selective control than the first one; actually, the base in the second position of the codon, for how the genetic code is defined, is the most important base in determining the chemical-physical properties of the corresponding amino acid. Despite the great overlap among the distributions of the classes, the differences observed among the slopes of the linear regressions are significant (*p* − *value* < 0.001).

**Figure 4.**
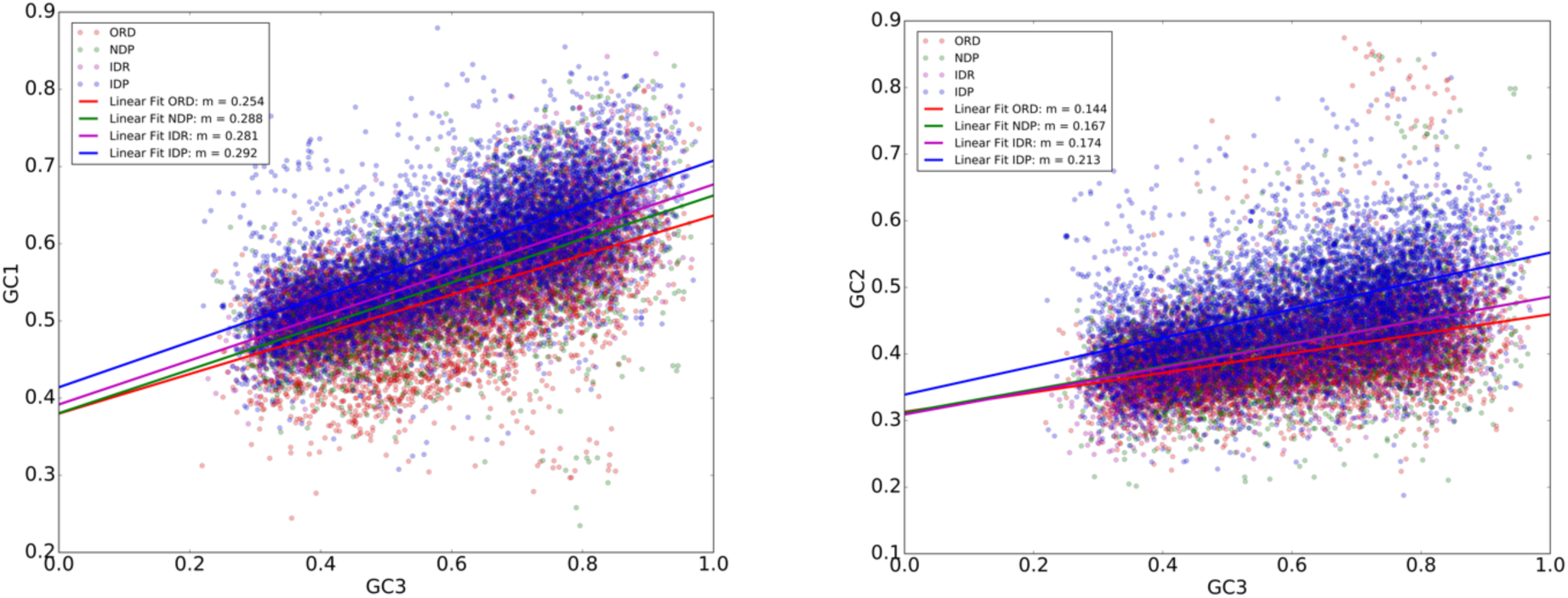
On the top, GC1 vs GC3. On the bottom, GC2 vs GC3. For each distribution, we do a linear fit and we report the slopes of the linear regressions in the legends.

In particular, the neutral selection acts more freely on all three codon positions in disordered variant (PDRs and IDPs) than in ordered ones (ORDs and NDPs). At the same time, from the slopes of the curves fit, we see that the third base of the codon has a much more marked influence with respect to the first two positions of codons, indicating the action of selective pressure on all variants of disorder.

### Variability in GC-content among variants of proteins

Since the purpose of this study is to analyze the evolutionary properties of the genes that encode the different proteins in our variants of disorder, another important parameter to take into account is the GC-content. In vertebrates, the GC-contents of genes are linearly related to those of the genomic regions in which they are located (Bernardi et al. 1985). GC-rich regions are gene-rich and high in several biological activities like transcription, translation, and recombination. It has been proposed that most housekeeping genes should be located in GC-rich isochores, whereas tissue-specific or developmentally regulated genes should be located in GC-poor isochores (Duret 2000). Many aspects of the isochores evolution in the human genome have long been investigated; nevertheless, it is not entirely clear whether GC-content is determined by neutral evolution or selection or both. For these reasons, it is very important to analyze the GC-content that characterizes our variants of disorder. In Fig. 5, we show the distributions of the GC-contents of the genes in each variants of disorder. From the form of distributions (unimodal for ORDs with a pick on the left toward low GC-content, similar bimodal distributions with two picks for NDPs and IDRs, and unimodal with one pick on the right toward high values of GC-content for IDPs) and the localization of the average values (green lines), passing from the ordered classes (ORDs and NDPs) to the disordered ones (PDRs and IDPs), we observed a shift of the average value towards higher values of the GC-content (Fig. 5). The difference observed in the average value is statistically significant (Mann-Whitney test with a *p* − *value* < 0.001).

**Figure 5.**
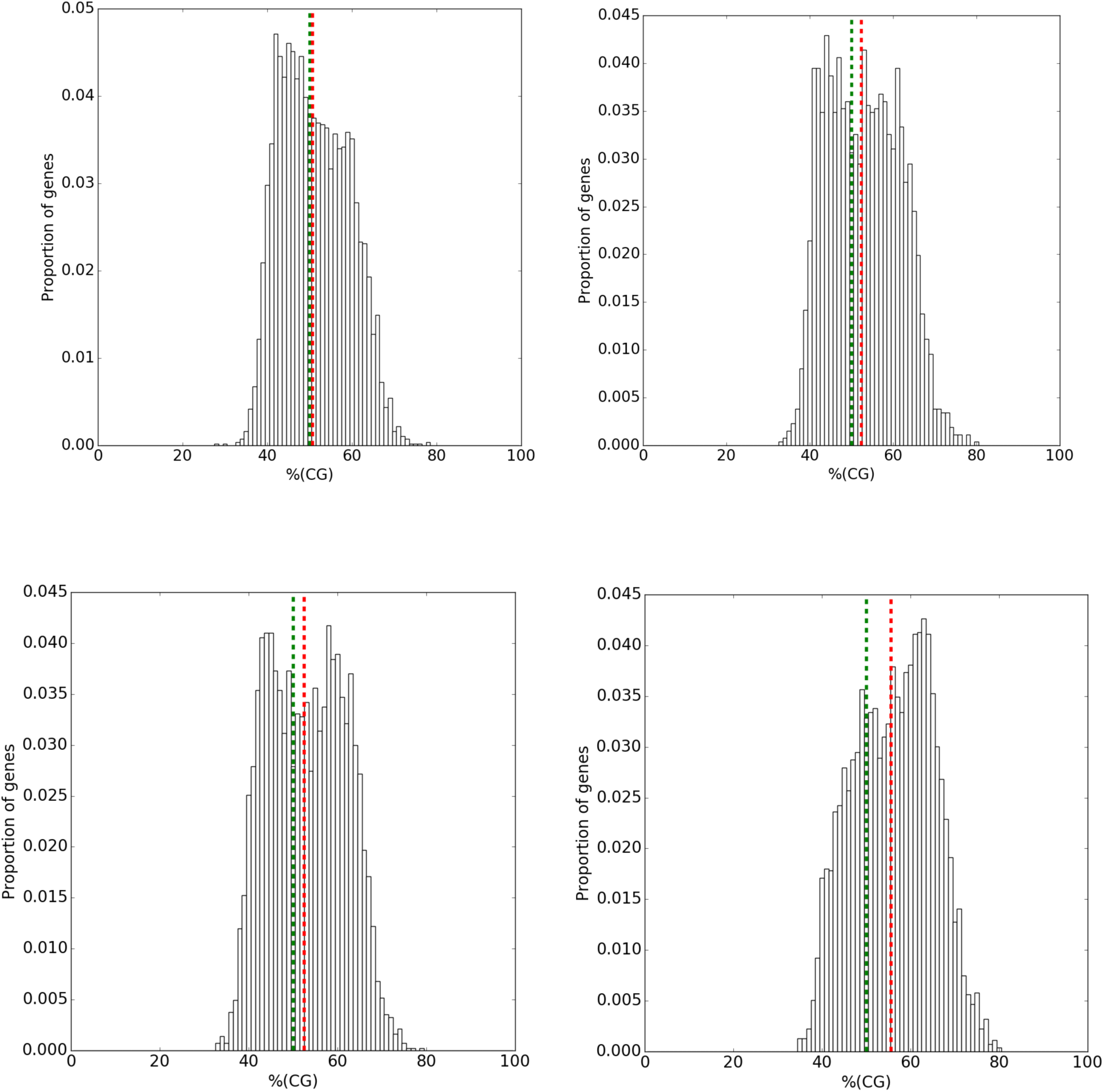
GC-content distributions for ORDs (top-left), NDPs (top-right), PDRs (bottom-left), and IDPs (bottom-right). The vertical red lines represent the average values of the distributions while the vertical green lines are located in corresponding of *GC* − *content* = 0.5.

Plotting the distributions of the TA-content for each class, we obtain exactly the specular behavior of the one just shown. So, increasing the percentage of disordered residues, the total GC-content of the corresponding gene increases (or decrease the percentage of the total TA-content of the corresponding gene). In Fig. 6, the average GC-content measured for each variant of disorder and for different percentiles of disordered residues in IDPs is shown; in line with Peng et al. (Peng et al. 2016), a monotonous trend is observed confirming the linear positive correlation between GC-content of the gene and the percentage of disordered residues in the corresponding amino acid sequence.

**Figure 6.**
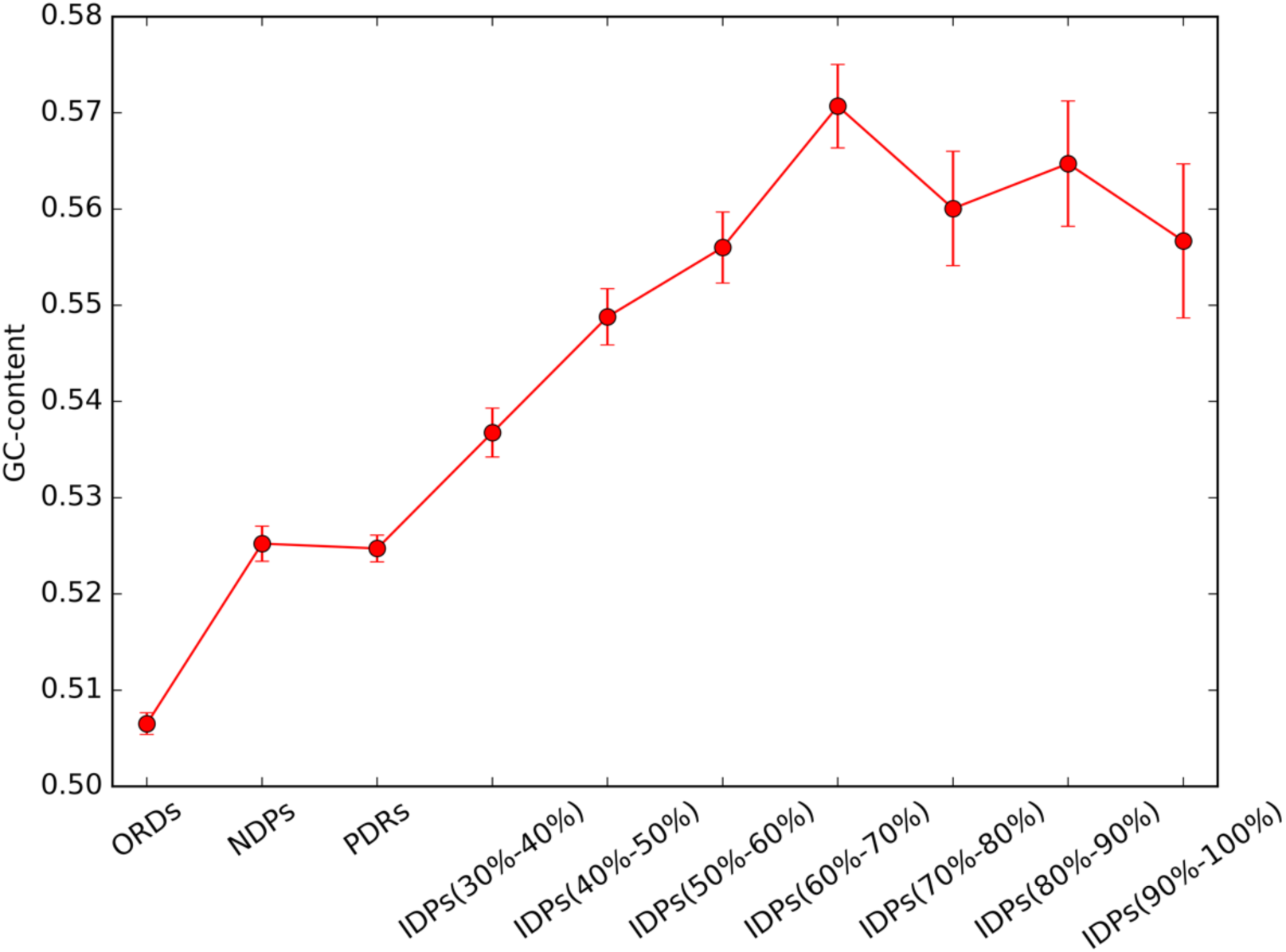
GC-content of the genes as a function of the percentage of disordered residues in the corresponding amino acid sequences. A positive monotonous trend is observed.

### Excluding compositional bias from the calculation of the selective pressure

Nucleotide and dinucleotide bias deeply influence CUB. There are three dinucleotide positions in a codon. Variation of dinucleotide 1–2 is strongly restricted by amino acid sequence; conversely dinucleotides 2-3 and 3-1 are more evolutionary flexible because they both include the variable third codon position. A general rationale to analyze the impact of (di)nucleotide bias on ENC is shuffling synonymous codons in the original genome sequence whilst preserving the factor under study (Belalov 2013). When shuffled sequences have an ENC equal to the original sequence, the constraint (e.g. the nucleotide content) fully explains the CUB (or, in simple words, at that (di)nucleotide composition ENC cannot possibly be higher than it is). On the other hand, a difference between ENC in the original and simulated sequences shows the extent of CUB that could not be explained by the kind of pressure (nucleotide or dinucleotide) that was used as a constraint for the shuffling algorithm (Belalov 2013). In Fig. 7, we show the ENC values for natural coding sequence and sequence obtained performing the various shuffling techniques. In sequences shuffled with preserved third codon position GC content (N3), ENC values were higher than the original ENC by more than 3 in all variants of disorder, indicating residual unexplained bias, but at the same time were below 61, indicating the effect of GC content on CUB. The impact of dinucleotide content on CUB was next tested. A notable disparity of dinucleotide frequencies at codon positions 2-3 and 3-1 (Fig. 7) justified treating them independently in shuffling algorithms. In all variants, shuffling of codon position 2-3 dinucleotide (dN_23_) produced ENC values that were lower by more than 4 than ENC upon N_3_ correction (Fig. 7) indicating a strong impact of position 2-3 dinucleotide content on CUB. Shuffling of dinucleotide position 3-1, on the contrary, did not result in a decrease of ENC, which differed from N_3_ shuffled sequences by less than 0.1/0.2 in all variants of disorder. To evaluate the impact of distinct dinucleotide content in codon positions 2-3 and 3-1 simultaneously, the codon position 2-3-1 triplet was shuffled between compatible codon pairs (dN_231_ correction). Consistent with results of dN_23_ and dN_31_ shuffling, ENC upon dN_231_ shuffling did not differ significantly from dN_23_ shuffling, supporting the negligible impact of dinucleotide content at codon positions 3-1 on synonymous codon preference.

**Figure 7.**
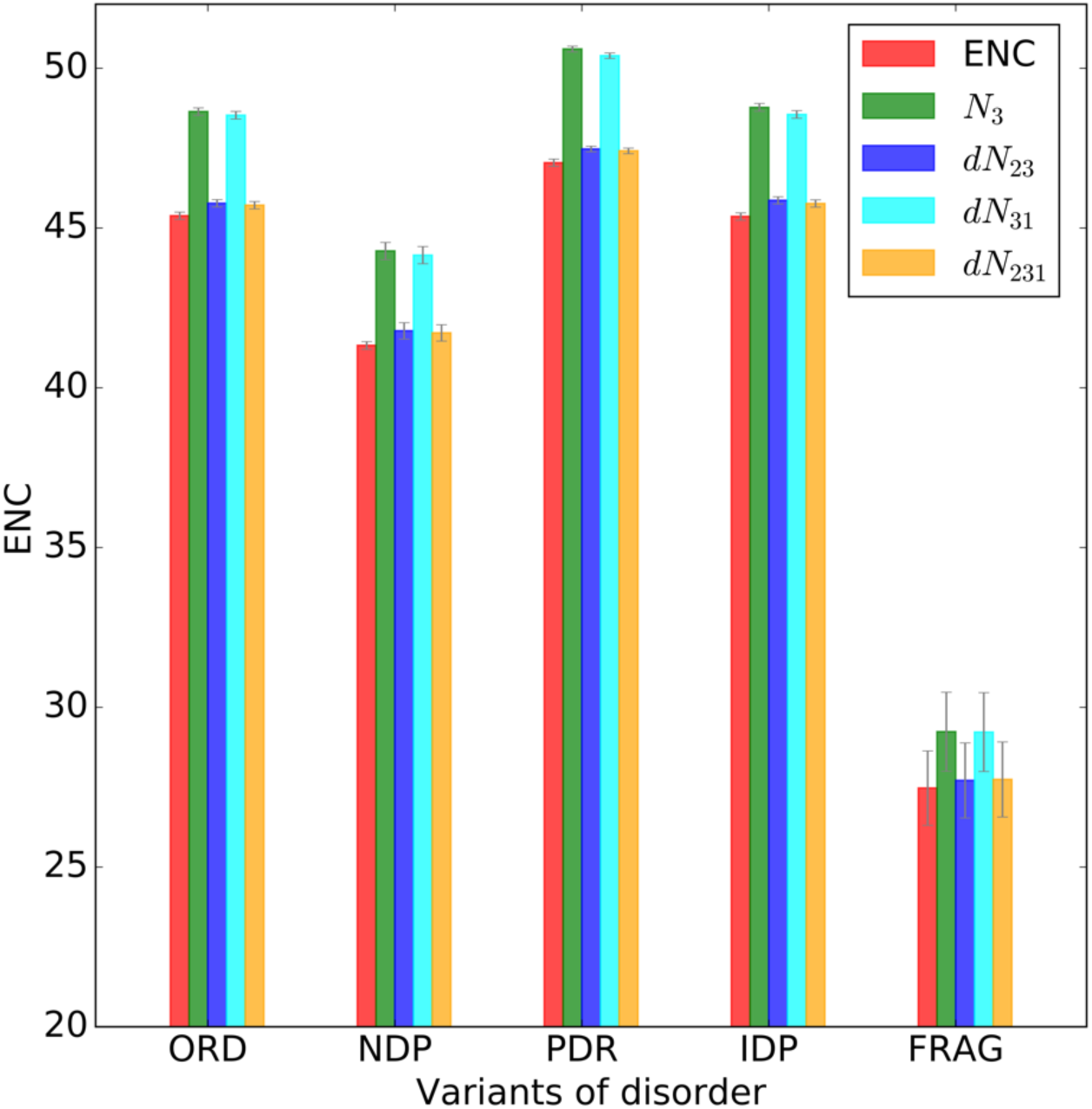
Average ENC values for the natural coding sequences (in red) and for shuffling sequences (other color) obtained in all variants of disorder.

It is worth noting that the differences obtained between the ENC values of shuffled sequences and ENC value of natural coding sequences allow us to quantify the extent of a notable residual unexplained bias. So, in Fig. 8 we report the differences obtained for all four shuffling techniques compared to the reference value obtained with the natural coding sequences. In this plot, closer to 0.0 is the average value of the difference and more we are justified in saying that the kind of pressure (nucleotide or dinucleotide) that was used as a constraint for the shuffling algorithm could be the cause of the codon bias.

**Figure 8.**
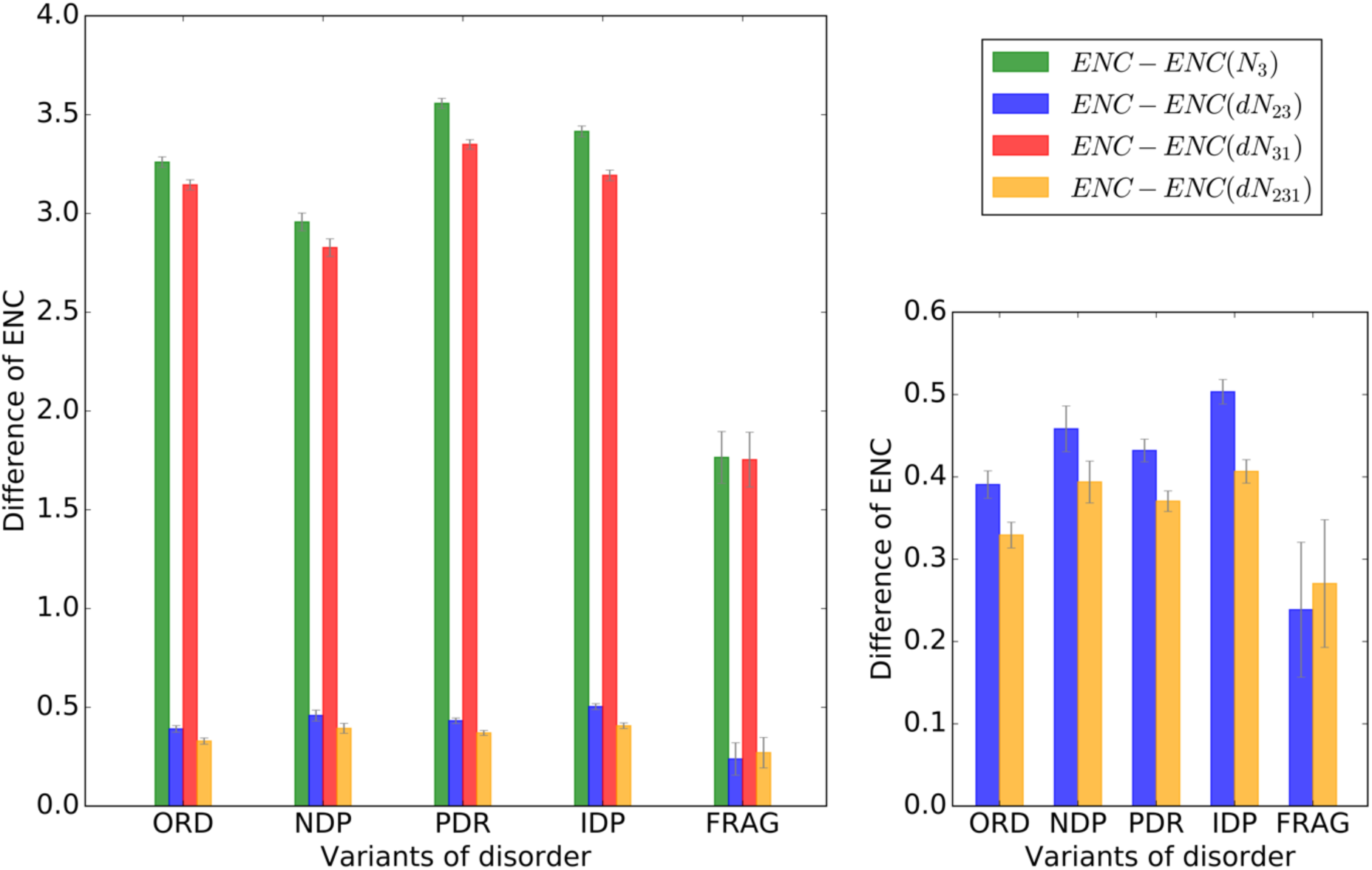
Average differences between ENC values obtained for N3, N23, N31, and N231 shuffled sequences and the corresponding ENC values of natural coding sequences. For greater clarify, on the bottom-right we report the average differences in ENC values obtained for N23 and N231 shuffled sequences and those for the original coding sequences.

Clearly, most of CUB could be attributed to mutational pressure on dinucleotide bias at codon position 2-3, which indicated the key role of mutational bias, but left notable unexplained bias (due to selective and/or translational pressure). We could say that the difference between the average values of ENC obtained performing a shuffling of dinucleotides at codon position 2-3 appears to be the most appropriate to quantify the effect of natural selection. To be more clear, in the insert on the bottom-right in Fig. 8 we show only the differences observed with N23 and N231 shuffling techniques. Notably, IDPs seem to be subject to a stronger residual unexplained bias identifiable with the action of natural selection. It is important to underline that even though the differences observed between IDPs and the other variants of disorder in terms of this residual unexplained bias are small (as expected), they are statistically significant (Matt-Whitney Test, p-value<0.00001).

### Fraction of CpG sites in the coding sequence and Transcription-Associated Mutational Biases (TAMB)

The CpG sites are regions of DNA where a cytosine nucleotide is followed by a guanine nucleotide in the linear sequence of bases along its 5' → 3' direction. Cytosines in CpG dinucleotides are mutational hotspots in mammalian genomes and they can be methylated to form 5-methylcytosine. In mammals, methylating the cytosine within a gene can change its expression, a mechanism that is part of a larger field of science studying gene regulation that is called epigenetics. Enzymes that add a methyl group are called DNA methyltransferases. The CpG notation is used to distinguish this single-stranded linear sequence from the CG base-pairing of cytosine and guanine for double-stranded sequences. CpG should not be confused with GpC, the latter meaning that a guanine is followed by a cytosine in the 5' → 3' direction of a single-stranded sequence. In the following, we report an example of CpG sites in the coding sequence.

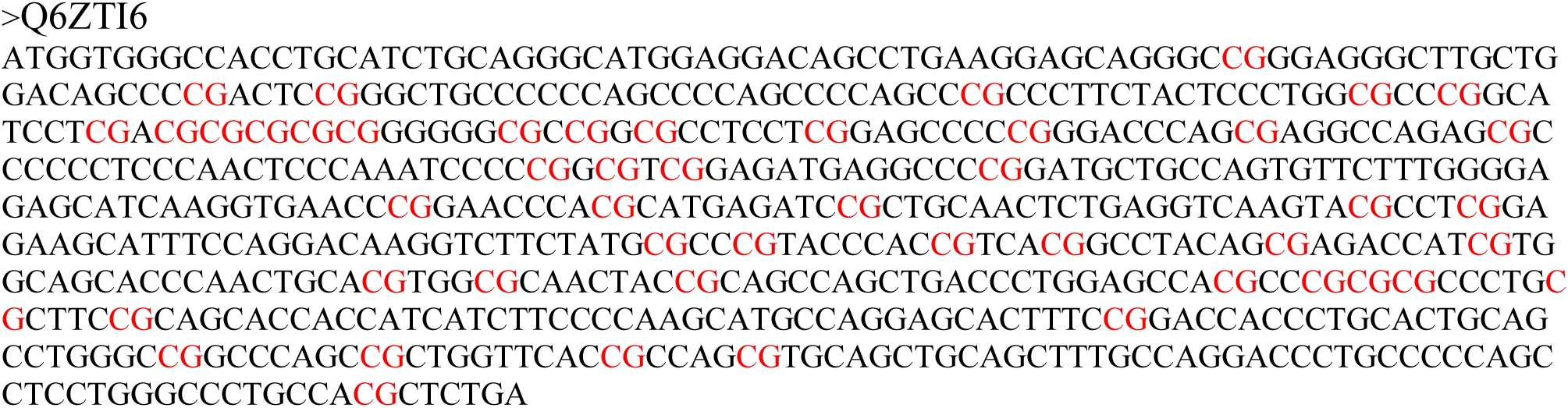

Our variants of disorder appear to be different in the fraction of sequence occupied by CpG sites. Interestingly, IDPs and FRAGs, which have a higher percentage of disordered residues, are characterized by a higher fraction of CpG sites in the sequence (Fig. 9). On the contrary, completely ordered proteins (ORDs) have the lowest fraction of CpG sites in the sequences (Fig. 9).

**Figure 9.**
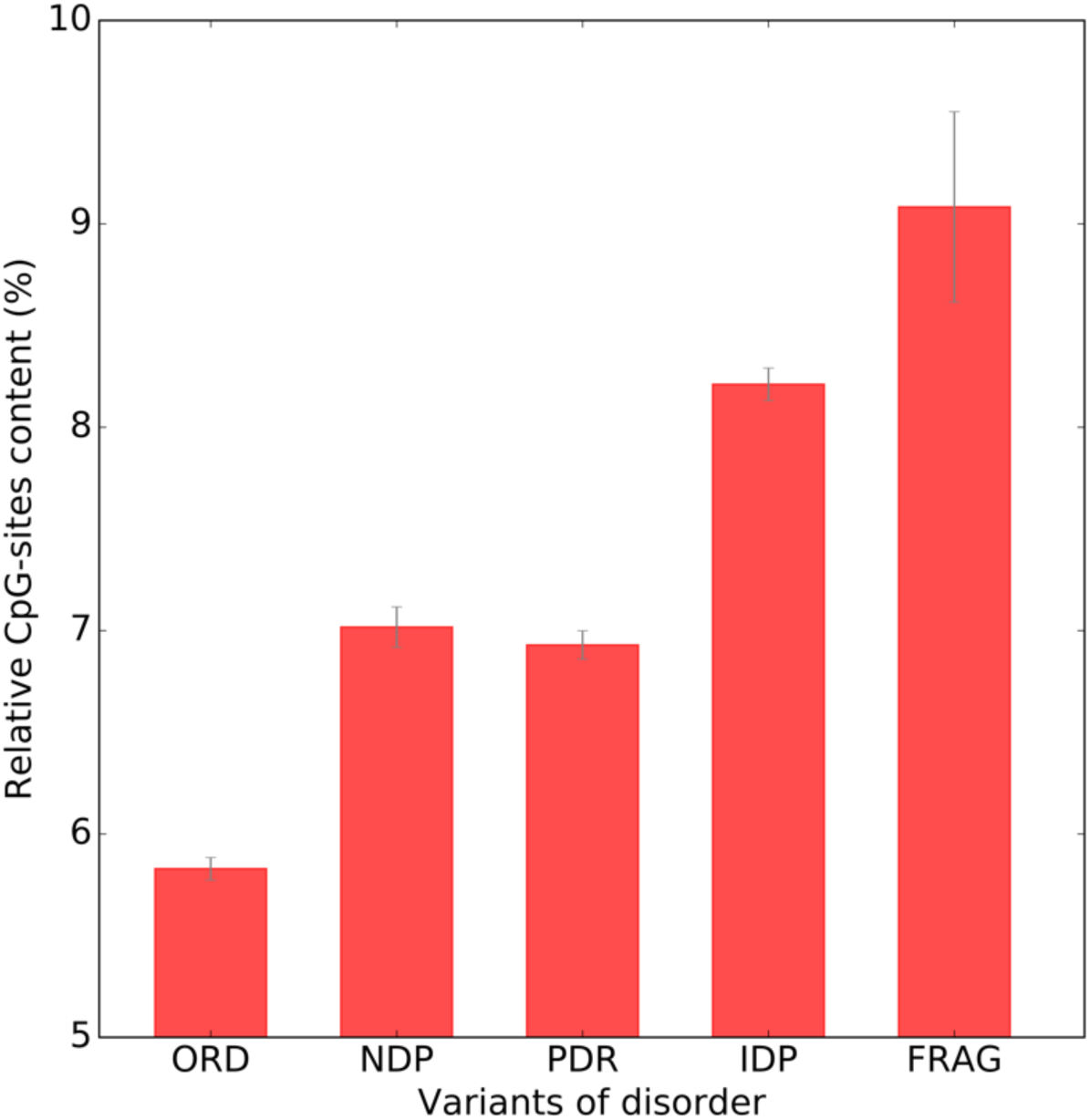
Average relative CpG-sites content in coding sequences in all variants of disorder.

These observations are quite interesting because many phenomena related to CpG sites are involved in cancers. Just think that the role of hypermethylation/hypomethylation of CpG island has been identified in almost all types of cancer (e.g. colon, stomach, pancreas, liver, kidney, lung, head and neck, breast, ovary, endometrium, kidney, bladder, brain, leukemia and lymphomas) (Esteller et al. 2001). Moreover, many cellular pathways are inactivated by this type of epigenetic lesion: DNA repair, cell cycle, apoptosis, cell adherence (Esteller 2002), confirming the central role of hypermethylation/hypomethylation of CpG island to progression to cancer. Coherently, we observed that cancer-related proteins are enriched in IDPs providing new insight about the role of this variant of disorder in human diseases (Deiana A., Forcelloni S., Porrello A., Giansanti A., pers. comm.).

Another aspect to take into account in a complex organism (like human) is the transcription-associated mutational biases (TAMB), which is expected to cause strand asymmetries, increasing G relative to C and T relative to A content of the coding strand (Comeron 2004). So, we investigated the consequences of TAMB by measuring G+T content in the coding sequences in each variant of disordered proteins. As shown in Figure 10, genes that encode for proteins in structured variants (ORDs, NDPs, PDRs) show the greatest influence of G+T content, evidencing TAMB. Conversely, genes encoding for proteins which are natively unfolded (IDPs) are characterized by the weakest effect of TAMB. Note that IDPs which are affected by the largest unexplained bias is the same variant showing the weakest TAMB (Fig. 10). To conclude this section, it is important to say that genes affected by TAMB will be subject to conflicting mutational and selective pressures on synonymous composition beyond the isochore effects. As a result, variants showing a larger effect of TAMB also reveal a minor influence of selection (unexplained bias) on synonymous codon usage. Conversely, variants showing less marked evidence of TAMB are those in which the extent of selection (unexplained bias) on synonymous composition appears to be larger (Fig. 10).

**Figure 10.**
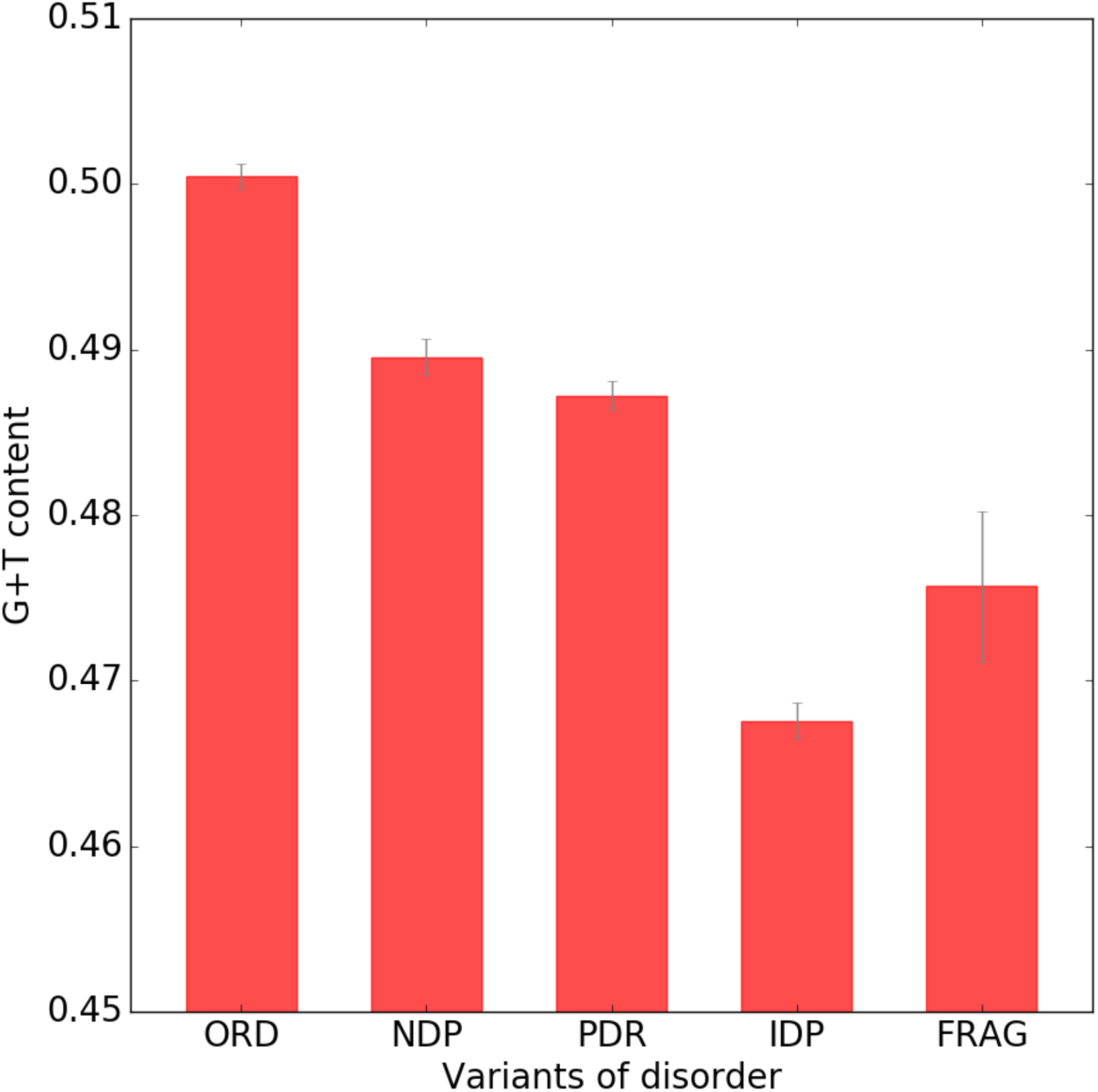
Average value of G+T content in the coding sequence in each variant of disorder as a measure of strand asymmetry.

## Discussion

A quantitative measure of the selective pressure acting on different proteins, especially in the human proteome, is very important in biotechnology and in many other fields of research. We need only think of heterologous gene expression, where the aim is to insert and make efficient the expression of a gene, or a part of a gene, in an organism where usually that gene does not occur. Clearly, the whole process has to be under control in order to minimize the risk of selecting and accumulating mutant variants of the corresponding protein that could destroy the cellular physiology of the host organism. A widely used method for avoiding deleterious effects on the fitness consists in optimizing the codon usage bias to that one of highly expressed genes in the host organism. Thus, knowing the various selective pressures acting on the genes in a given organism could have a positive the heterologous gene expression minimizing deleterious changes for the physiologic state but, at the same time, maximizing gene expression.

In this work, we perform a systematic analysis of the mutational bias and selective pressure acting in the human proteome, using the codon usage bias to assess the evolutionary pressures driving the long time scales genome evolution. In doing this, we separate the human proteome in five variants of disorder (ORDs, NDPs, PDRs, IDPs, and FRAGs) classified on the basis of the type of predicted disorder in the sequence. It is important to point out that the signals we are looking for if it exists at all, is expected to be very weak, so that it may be detected only in very large sequence ensembles. In the first two sections, using different genetic tools, such as ENC-plot, PR2-plot, Neutrality plot, and GC-content, we concluded that proteins containing long disordered regions accommodated in structured domains (PDRs) and proteins with high percentage of disordered residues in the sequences (IDPs) are more affected by the neutral changes of their sequences, which probably tend to stabilize thermodynamically the genetic material (see distributions of average thermostability of codons that compose the gene sequences in Supplementary Material – Fig. S1). To put it another way, these observations mean that evolutionary rates in PDR and IDP sequences are on average higher with respect to ORD and NDP sequences. So, PDR and IDP sequences diverge more rapidly than ORDs and NDPs sequences. Probably this characteristic of IDPs is due to the fact that in structured or globular proteins few mutations (even between synonymous codons) in the corresponding gene, which is a section of DNA that provides the blueprint for a specific protein by regulating the velocity of the translational process, can result in a protein that no longer folds properly. That makes the protein useless because it is the shape of a protein that largely determines which molecules it can bind to, and hence, its function. On the contrary, in specific unstructured regions of PDRs and especially in IDP variant, structural constraints are generally fewer and often of a more local nature, making them more prone to accepting neutral mutations. To confirm this, just think that in particular subclasses of PDRs and IDPs, such as entropic chains, may be under only two very general constraints: keeping the region disordered and its length unchanged. Nevertheless, this does not mean that IDPs are not affected by selective pressure. Actually, through this analysis, we found that IDPs are also those most subject to an unexplainable pressure identifiable with some selective mechanism that differentiates them from the others in a statistically significant way.

Taken together, our results show that there are different levels of evolutionary pressure acting in the human proteome and that could be useful considering them in future studies about synthetic biology and biotechnology applications. Recently, Afanasyeva et al. suggest that disordered protein regions are important targets of genetic innovation and that the contribution of positive selection in these regions is more pronounced than in other protein parts (Afanasyeva et al. 2018). Coherently with these observations, through an independent procedure based only on codon statistics in coding sequences, we underline the importance of the unstructured regions and natively unfolded proteins from and evolutionary standpoint being freer to accept mutations, both neutral and selective.

## Supporting information

