## Supplementary material for "New perspectives into the evolutionary pressures acting on the Human intrinsically disordered proteins"

Supplementary Informations

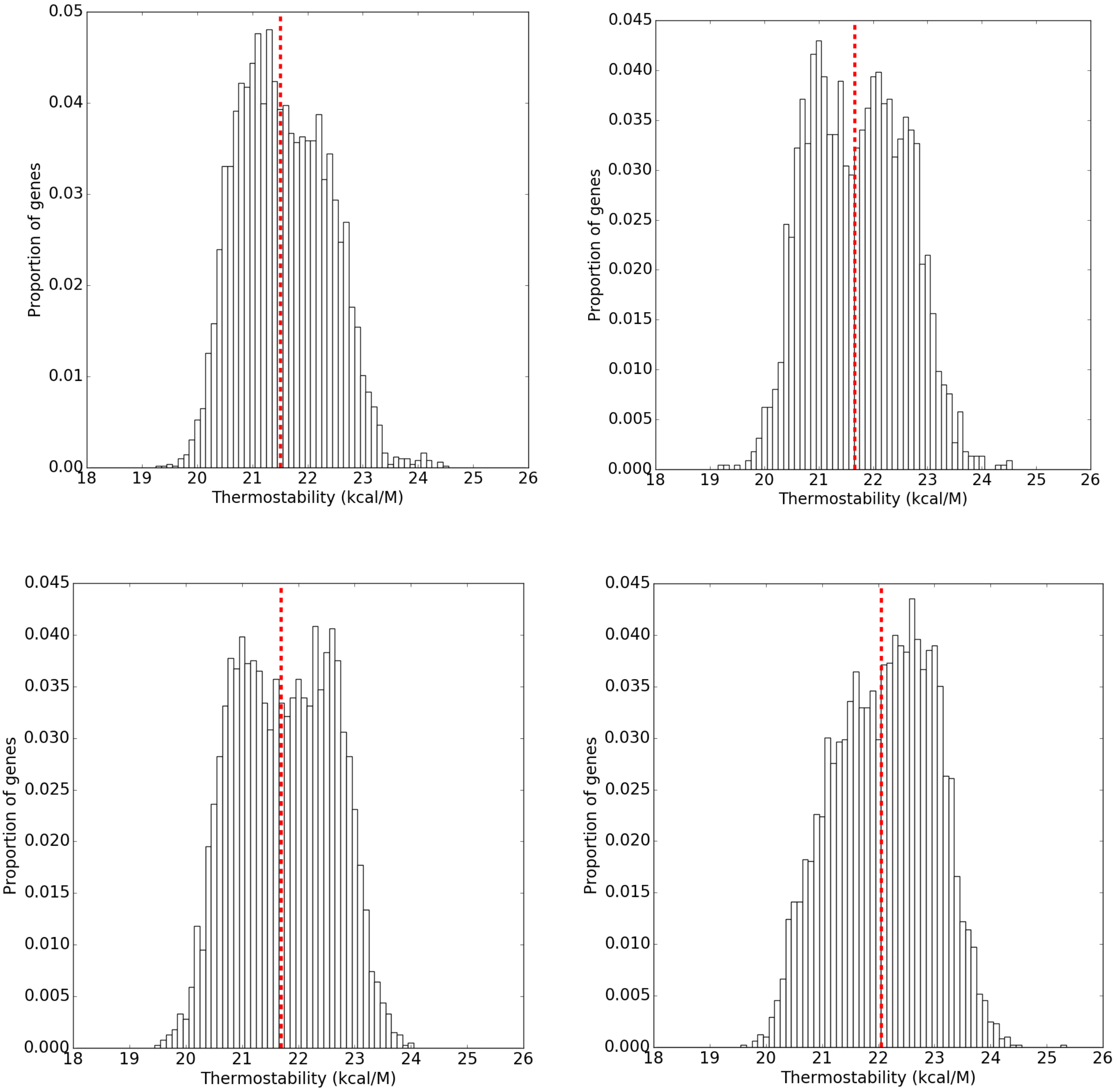


**Figure S1**. Distributions of the average thermostability of codons that compose the gene sequences of ORDs (top-left), NDPs (top-right), PDRs (bottom-left), and IDPs (bottom-right). The vertical red lines represent the average values of the distributions.
